# Adaptive introgression from distant Caribbean islands contributed to the diversification of a microendemic radiation of trophic specialist pupfishes

**DOI:** 10.1101/115055

**Authors:** Emilie J. Richards, Christopher H. Martin

**Affiliations:** Biology Department, University of North Carolina at Chapel Hill, Chapel Hill, North Carolina, United States of America

**Keywords:** adaptive introgression, speciation, adaptive radiation, hybridization

## Abstract

Rapid diversification often involves complex histories of gene flow that leave variable and conflicting signatures of evolutionary relatedness across the genome. Identifying the extent and source of variation in these evolutionary relationships can provide insight into the evolutionary mechanisms involved in rapid radiations. Here we compare the discordant evolutionary relationships associated with species phenotypes across 42 whole genomes from a sympatric adaptive radiation of *Cyprinodon* pupfishes endemic to San Salvador Island, Bahamas and several outgroup pupfish species in order to understand the rarity of these trophic specialists within the larger radiation of *Cyprinodon.* 82% of the genome depicts close evolutionary relationships among the San Salvador Island species reflecting their geographic proximity, but the vast majority of the fixed variants between the specialist species lie in regions with discordant topologies. These regions include signatures of selective sweeps and adaptive introgression from neighboring islands into each of the specialist species. Hard selective sweeps of genetic variation on San Salvador contributed 10-fold more to divergence between specialist species within the radiation than adaptive introgression of Caribbean genetic variation; however, some of these introgressed regions from distant islands were associated with the primary axis of oral jaw divergence within the radiation. For example, standing variation in a proto-oncogene (*ski*) known to have effects on jaw size introgressed into one San Salvador specialist from an island 300 km away. The complex emerging picture of the origins of adaptive radiation on San Salvador indicates that multiple sources of genetic variation contributed to the adaptive phenotypes of novel trophic specialists on the island. Our findings suggest that a suite of factors, including rare adaptive introgression, may also be required to trigger adaptive radiation in the presence of ecological opportunity.

**Author summary:** Groups of closely related species can rapidly evolve to occupy diverse ecological roles, but the ecological and genetic conditions that trigger this diversification are still highly debated. We examine patterns of molecular evolution across the genomes of a rapid radiation of pupfishes that includes two trophic specialists. Despite apparently widespread ecological opportunities and gene flow across the Caribbean, this radiation is endemic to a single Bahamian Island. Using the whole genomes of 42 pupfish we find evidence of extensive and previously unexpected variation in evolutionary relatedness among Caribbean pupfish. Two sources of genetic variation have contributed to the adaptive diversification of complex phenotypes in this system: selective sweeps of genetic variation from across the Caribbean that was brought into San Salvador through hybridization and genetic variation found on San Salvador. While genetic variation from San Salvador appears to be relatively more common in the divergence observed among specialists, hybridization probably played an important role in the evolution of the complex phenotypes as well. Our findings that multiple sources of genetic variation contribute to the San Salvador radiation suggest that a complex suite of factors, including hybridization with other species, may be required to trigger adaptive radiation in the presence of ecological opportunity.

## Introduction

Adaptive radiations are central to our understanding of evolution because they generate a wealth of ecological, phenotypic, and species diversity, often in very rapid bursts. However, the mechanisms that trigger the rapid bursts of trait divergence, niche evolution, and diversification characteristic of classic adaptive radiations are still debated. The availability of resources in new environments with few competitors has long been seen as the major force driving adaptive radiations[1–3], but it is a longstanding question why only some lineages rapidly diversify in response to such resource abundance while others do not [4–9].

Gene flow can introduce adaptive genetic variants [10,11], genetic incompatibilities [12–14], and/or reinforcements [15–18] that initiate or contribute to the process of speciation and a growing number of studies have identified gene flow and genome-wide introgression across a range of adaptive radiations [19–26]. The hybrid swarm hypothesis [27] proposes that hybridization among distinct lineages can introduce genetic diversity and novel allele combinations genome-wide that may trigger rapid diversification in the presences of abundant ecological opportunity. Yet there is still little evidence that hybridization specifically triggered adaptive diversification within radiations, as opposed to simply being pervasive throughout the history of any young rapidly diversifying group [25,28]. One of the only examples with strong evidence of hybridization leading to ecological and species diversification is that of several hybrid species within a radiation of *Helianthus* sunflowers [29–34]. However, this may simply represent examples of multiple homoploid speciation events within an already radiating lineage rather than a hybrid swarm scenario. So while there is convincing evidence that hybridization can facilitate diversification among species pairs (but see [26,35] for a potential multispecies outcome of hybridization), it is still unclear whether gene flow is a major factor constraining adaptive radiation in some lineages or if ecological opportunity is the sole constraint.

The adaptive radiation of San Salvador pupfishes provides an outstanding system to address these questions about the contributions of different sources of genetic variation to rapid diversification and the role of gene flow in the evolution of complex phenotypes. Pupfish species of the genus *Cyprinodon* inhabit saline lakes and coastal areas across the Caribbean and Atlantic and nearly all pupfishes are dietary generalists consuming algae and small invertebrates [36]. In contrast, three *Cyprinodon* species live sympatrically in the hypersaline lakes of San Salvador Island and comprise a small radiation that has occurred within the past 10,000 years based on the most recent glacial maximum when these lakes were dry due to lowered sea levels [37–39]. This radiation is composed of the widespread generalist algae-eating species *Cyprinodon variegatus* and two endemic specialists that coexist with the generalist in all habitats in some lakes. These specialists have adapted to unique trophic niches using novel morphologies: the molluscivore *Cyprinodon brontotheroides* with a unique nasal protrusion and the scale-eating *Cyprinodon desquamator* with enlarged oral jaws and adductor mandibulae muscles [36,40]. Surveys of populations living on neighboring islands in the Bahamas and phylogenetic analyses with other *Cyprinodon* species indicate that these specialist species are endemic to the hypersaline lakes of San Salvador Island and that both specialists arose from a generalist common ancestor during this recent radiation [41].

The currently available ecological and genetic data on the group provides little indication as to why this radiation is localized to a single island. Variation in ecological opportunity among hypersaline lake environments in the Caribbean does not appear to explain the rarity of this radiation [41]. This finding suggests a potentially important role for sufficient genetic variation to respond to abundant, underutilized resources in these environments. However, a hybrid swarm hypothesis about the origins of the radiation does not appear to explain its rarity either: genetic diversity is comparable among islands and gene flow occurs among all Caribbean islands investigated, not only into San Salvador Island [41]. Novel traits and increased rates of diversification associated with them are well documented in this system [36,41,42], but understanding the rarity of this adaptive radiation requires a thorough investigation of the underlying genetic variation that accompanies these rare ecological transitions. A recent study investigating the genetic basis of trophic specialists in this radiation revealed very few regions underlying these phenotypes [43]. Only hundreds of variants out of 12 million were fixed within the scale-eater and molluscivore species. Since genetic divergence is limited to particular regions, localized rather than genome-wide investigations of the genome will be important for understanding how genetic variation, possibly originating outside of San Salvador, has contributed to the exceptional phenotypic diversification restricted to this island. Here, we use a machine-learning approach to identify regions of the genome with different evolutionary histories across 42 pupfish genomes sampled from the San Salvador radiation, two distant Caribbean islands, and 3 additional outgroups. We then scan the genome for evidence of localized introgression with pupfish populations outside of San Salvador Island and compare the relative contributions of adaptive introgression from two distant islands and hard selective sweeps to the divergence of each specialist species.

## Results

### Extensive variation in patterns of evolutionary relatedness across the genome

To identify localized patterns of population history across the genome, we used the machine-learning approach SAGUARO. SAGUARO combines a Hidden Markov Model with a self-organizing map to characterize local histories across the genome among aligned individuals [44]. This method does not require any *a priori* hypotheses about the relationships among individuals, but rather infers them directly from the genome by finding regions of consecutive nucleotides with a similar pattern of genetic differentiation, building hypotheses about relationships among individuals from these genetic differences, and then assigning regions of the genome to these hypothesized local “histories”. Since smaller segments with fewer informative SNPs are more likely to be incorrectly assigned to a hypothesized history by chance (pers. comm. M.G. Grabherr), we filtered out segments with fewer than 20 SNPs. Using this approach, we partitioned the genome into a total of 15 unique histories across 227,248 genomic segments that ranged from 101-324,088 base pairs in length (median: 852 bp) (S1 and S2 Figs Table S1).

The most prevalent history across 64% of the genome featured the expected species phylogeny for this group from previous genome-wide studies [36,41,45], in which all individuals from San Salvador Island were cleanly grouped into a single clade with distant relationships to generalist pupfish populations from other islands in the Caribbean, Death Valley in California, and two specialists from a second radiation in Mexico spanning the most divergent branch of the *Cyprinodon* tree (Fig 1). Unlike previous genome-wide phylogenies [41,45], and with the exception of a few individuals that grouped with molluscivores by lake, the generalists on San Salvador form a discrete clade from the molluscivores and scale-eaters.

**Fig 1.**
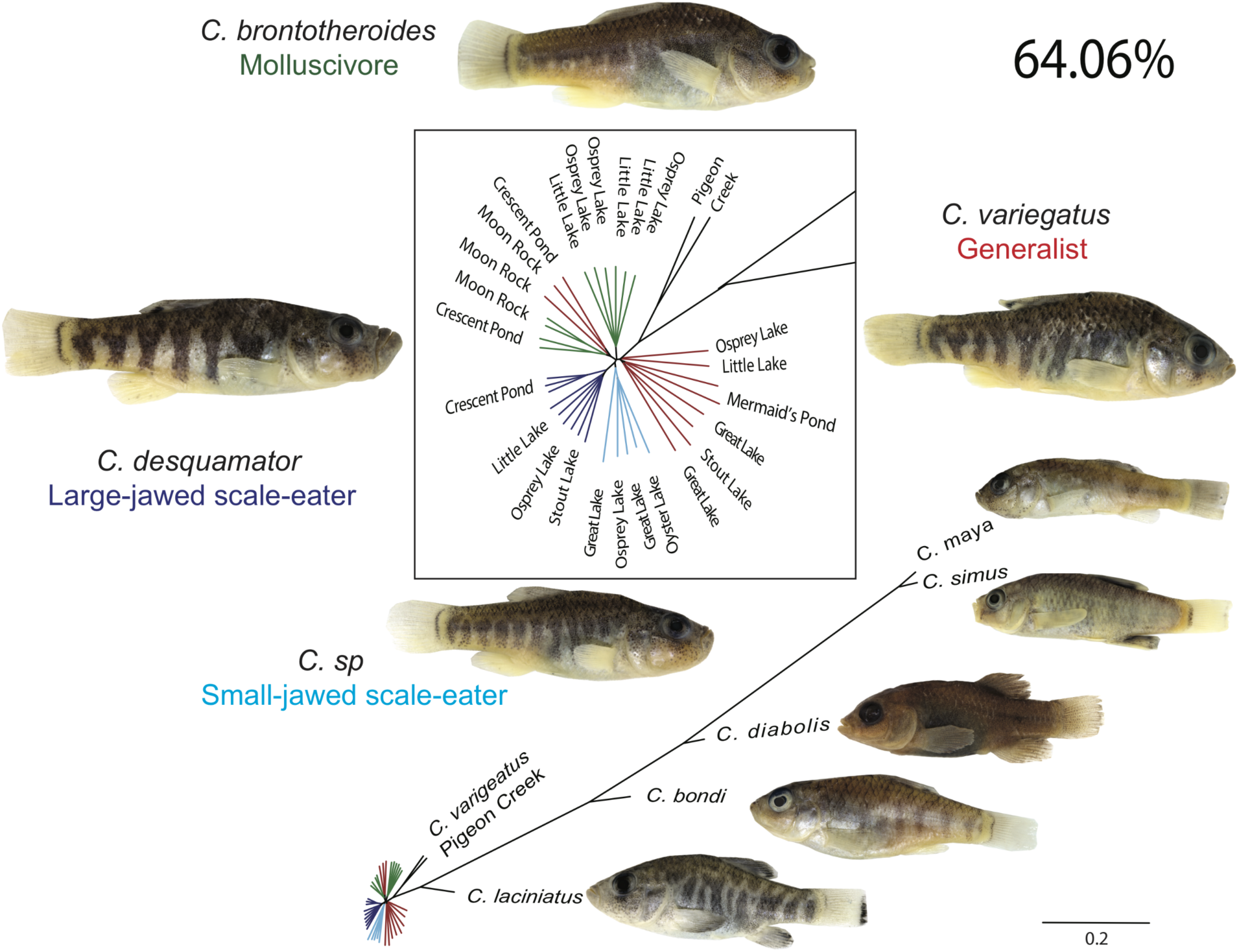
Neighborhood-joining tree of the dominant history of a monophyletic San Salvador clade covering 64% of the genome. San Salvador generalists (red), molluscivores (green), large-jawed scale-eaters (dark blue), small-jawed scale-eaters (light blue), and outgroup species (black) in the Caribbean, California, and Mexico. Other histories featuring a monophyletic San Salvador clade are presented in S1 Fig.

Within this dominant topology, scale-eaters from six lakes on San Salvador fell into one of two separate clades: small-jawed individuals from Osprey Lake, Great Lake, and Oyster Pond and larger-jawed individuals from Crescent Pond, Stout Lake, Osprey Lake, and Little Lake (Fig 1).

Molluscivores did not form a single clade as individuals from some lakes (Crescent Pond and Moon Rock) were more closely related to generalists from the same lake than molluscivores from other lakes, similar to previous genome-wide phylogenies [45]. Another history covering 10% of genome was very similar to the dominant one, differing only in the relationships among San Salvador generalists (S1 Fig). Additional histories spanning 7.6% of the genome featured a single San Salvador clade but also depicted a closer relationship between San Salvador and the outgroups as well as groupings of all three San Salvador species by lake in Crescent Pond and Moon Rock Pond. When combined with the dominant history, only 82.6% of the genome supported the expected San Salvador clade (S1 Table).

In other regions of the genome, San Salvador did not form a single clade (Figs 2A-C and S2, S1 Table). The most frequently observed alternative relationships depicted specialist individuals as a clade outside of the San Salvador Island group and sister to all the outgroup *Cyprinodon* species (Figs 2A,B). One such history, the ‘large-jawed scale-eater topology’, featured large-jawed scale-eater individuals outside of the San Salvador clade and was assigned to 4,437 segments covering 3.27% of the genome (Fig 2A). Another history, the ‘molluscivore topology’ showed a similar pattern in which the molluscivore individuals form a single clade outside of the San Salvador group and sister to all other outgroups (Fig 2B). This molluscivore topology was assigned to 3,916 segments and covered 3.11% of the genome. Another 2,029 segments covering 1.66% of the genome were assigned to a topology where both the large-jawed and small-jawed scale-eaters formed a combined clade outside of the San Salvador group, the “combined scale-eater topology’ (Fig 2C). Other histories featuring one of the specialists separated from the rest of San Salvador covered 0.76%-2.48% of the genome (Table S1).

**Fig 2.**
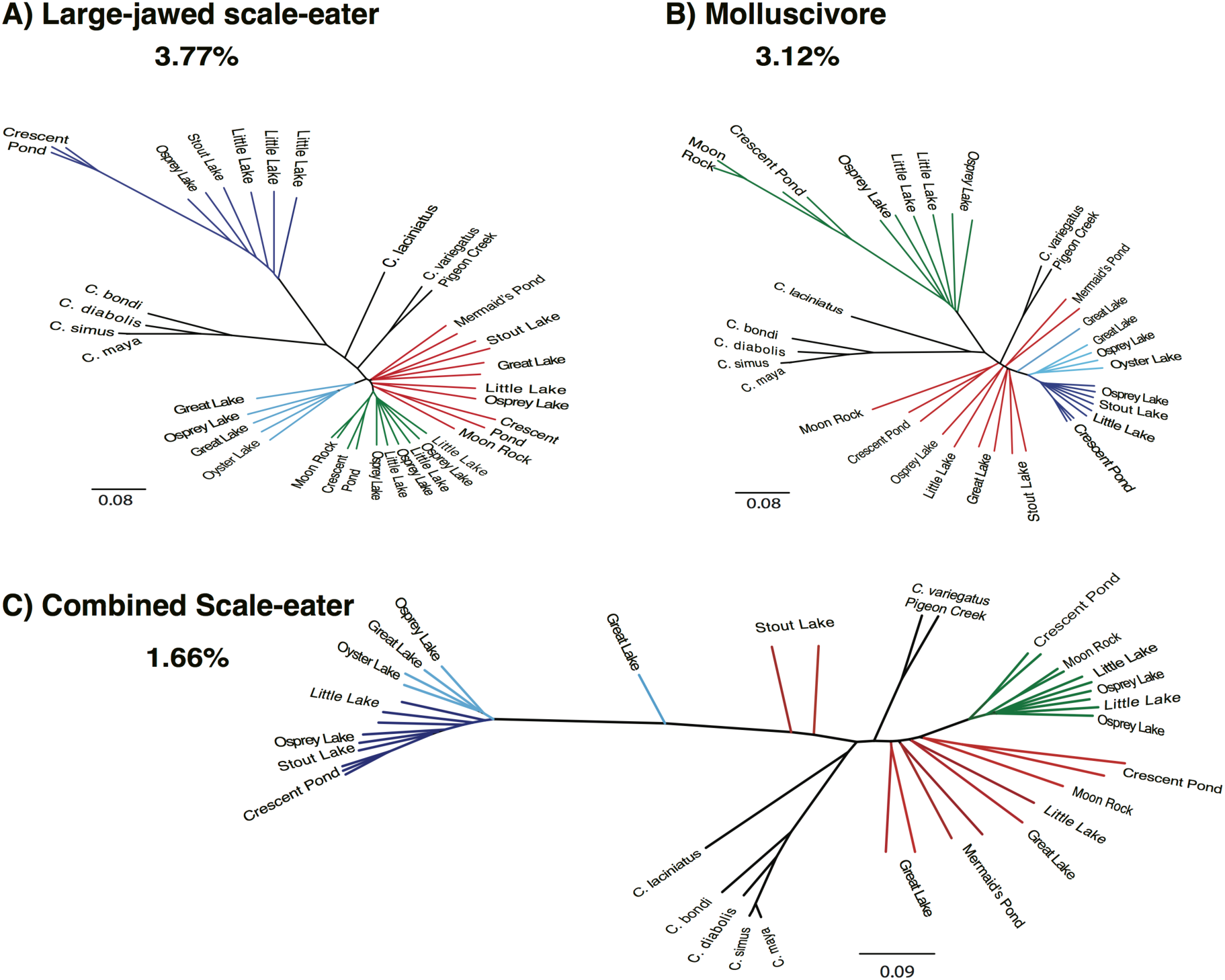
Neighborhood-joining trees of alternative topologies. (A) The large-jawed scale-eater topology describing 3.77% of the genome, where larger jawed scale-eater individuals fall outside of the San Salvador Island clade, with a sister relationship to outgroup pupfish species. (B) The mollsucivore topology assigned to a non-overlapping 3.12% of the genome, where molluscivore individuals fall outside of the San Salvador Island clade, with a sister relationship to the outgroup pupfish species. (C) The combined scale-eater topology assigned to another non-overlapping 1.66% of the genome, where all scale-eaters (along with two generalists from Stout’s Lake) fall outside of the San Salvador clade with a sister relationship to the outgroup pupfish species. Additional alternative topologies are presented in Fig S2.

Unexpectedly, all histories separated scale-eaters into groups of smaller- and larger-jawed individuals and the relationships between these two groups and other species differed across different regions of the genome. In some regions, the small-jawed scale-eater individuals were sister to the large-jawed scale-eaters (Figs 1,2B-C, and S1). In other regions, the small-jawed scale-eaters were more closely related to the generalists and molluscivores (Figs 2A and S1). These small-jawed scale-eaters may be a product of ongoing hybridization between species on San Salvador, an intermediate stage in the evolutionary path towards scale-eating, or a new species. Their consistent appearance as a separate clade supports a previously unrecognized small-jawed scale-eating species in some lakes.

### Localized introgression into both specialists from across the Caribbean

We examined signals of introgression from two distant pupfish generalist populations in the Caribbean: Lake Cunningham, New Providence Island in the Bahamas (described as the endemic species *Cyprinodon laciniatus* [46]) and Etang Saumautre Lake in the Dominican Republic (described as the endemic species *Cyprinodon bondi* [47]). We characterized the genomic landscape of introgression in the three San Salvador species using *f_4_* statistics that were initially developed to test for introgression among human populations [48–50].

We found 230 10-kb regions out of 100,260 that contained significant evidence of introgression between *C. laciniatus* or *C. bondi* and the San Salvador specialists (Fig 3A). Introgressed regions were scattered across the genome in 140 of the 9,259 scaffolds in our dataset. These regions were not typically concentrated in one section of the genome, with the largest cluster within a single scaffold containing only 5% of the total regions detected (Fig 3A). The genomic regions with significant evidence of introgression varied between the two specialists (Fig 3B,C), suggesting that admixture with other Caribbean populations has occurred multiple times and independently for each specialist or that different introgressed regions were used by the two specialists after a single admixture event (see S3-S5 Figs for full Manhattan plots). Only 23 regions of the 205 and 236 regions with significant evidence of introgression were shared between generalist/scale-eater and generalist/molluscivore comparisons, respectively. We also tested for introgression with the small-jawed scale-eaters excluded to search for potential introgression with the large-jawed scale-eaters alone (S6 Fig). Introgressed regions were less variable between the two groups of scale-eaters, with 150 of 211 regions shared. The sixty-one introgressed regions unique to the large-jawed scale-eaters suggest that some introgression occurred between populations on other Caribbean islands and the large-jawed scale-eater population, independently from the small-jawed scale-eaters.

**Fig 3.**
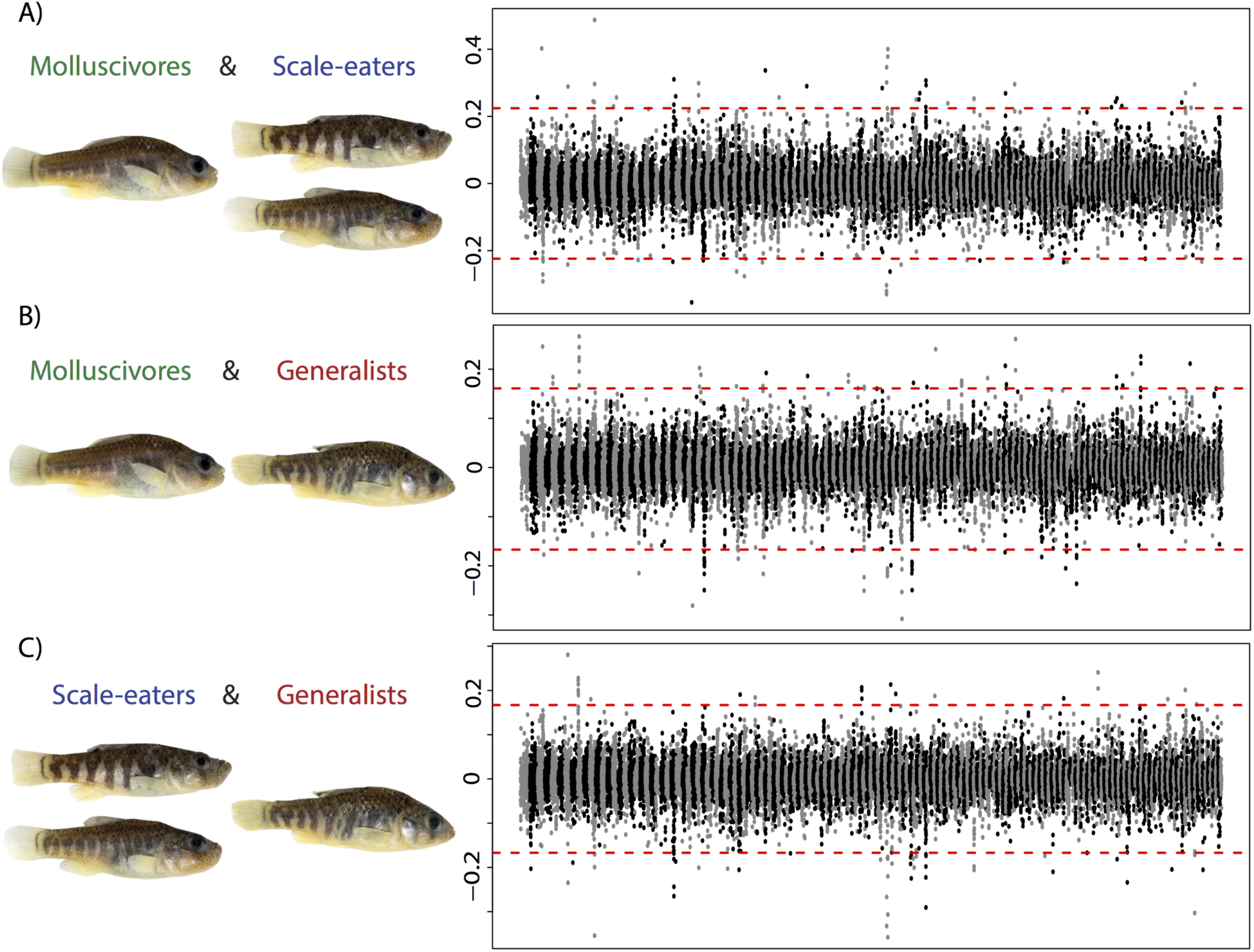
Variable introgression from distant Caribbean islands across the genomes of the San Salvador trophic specialists. Manhattan plot of the *f4* values between the *C. laciniatus* from New Providence Island, Bahamas, *C. bondi* from Etang Saumautre Dominican Republic and (A) molluscivores and scale-eaters on San Salvador, (B) molluscvires and generalists from San Salvador, (C) scale-eaters and generalists on San Salvador. Alternating gray/black colors indicate different scaffolds from the largest 170 scaffolds of the genome. Dotted red lines mark the of Bonferroni-corrected significance level for the *f*_4_ values (P-value < 5.2×10^−7^). Full Manhattan plots for each comparison are presented in Figs. S3-S6.

To assess whether our regions of introgression were predominately from regions of low diversity, we looked at *D_xy_* and π estimates across the detected regions of introgression in comparison to the genome-wide estimates (mean *D_xy_= 0.116*; mean π scale-eater=*0.0035*; mean π molluscivore *=0.0044*). We found that while some of the regions with significant *f_4_* values were in regions with low *D_xy_* and/or π between the San Salvador species, the majority were in regions of intermediate diversity (S7 Fig), and some were in regions with higher between-population divergence between the two specialists (S7 Fig), consistent with introgression that contributed to speciation.

### Multiple sources of genetic variation underlie species divergence

The relationships observed in the three alternative topologies (Fig 2) underlie most of the divergence observed between the molluscivores and scale-eaters, as 75% and 88% of the fixed SNPs between molluscivores and large-jawed scale-eaters and molluscivores and all scale-eaters respectively fall in these histories that make up less than 5% of the genome in total. Many of these regions contained candidate genes previously associated with variation in *Cyprinodon* jaw size [43]: 18 of the 31 candidate regions occurred in the combined scale-eater topology, and 1 candidate region in the molluscivore topology.

We then assessed the relative contributions of different sources of genetic variation to the divergence observed between the two specialists. The overlaps between these signals are summarized in Fig 4 (also see S8 Fig). The alternative topologies contained a greater proportion of regions with introgressed genetic variation and selective sweeps than those regions assigned to the dominant topology (Figs 4 and S8).

**Fig 4.**
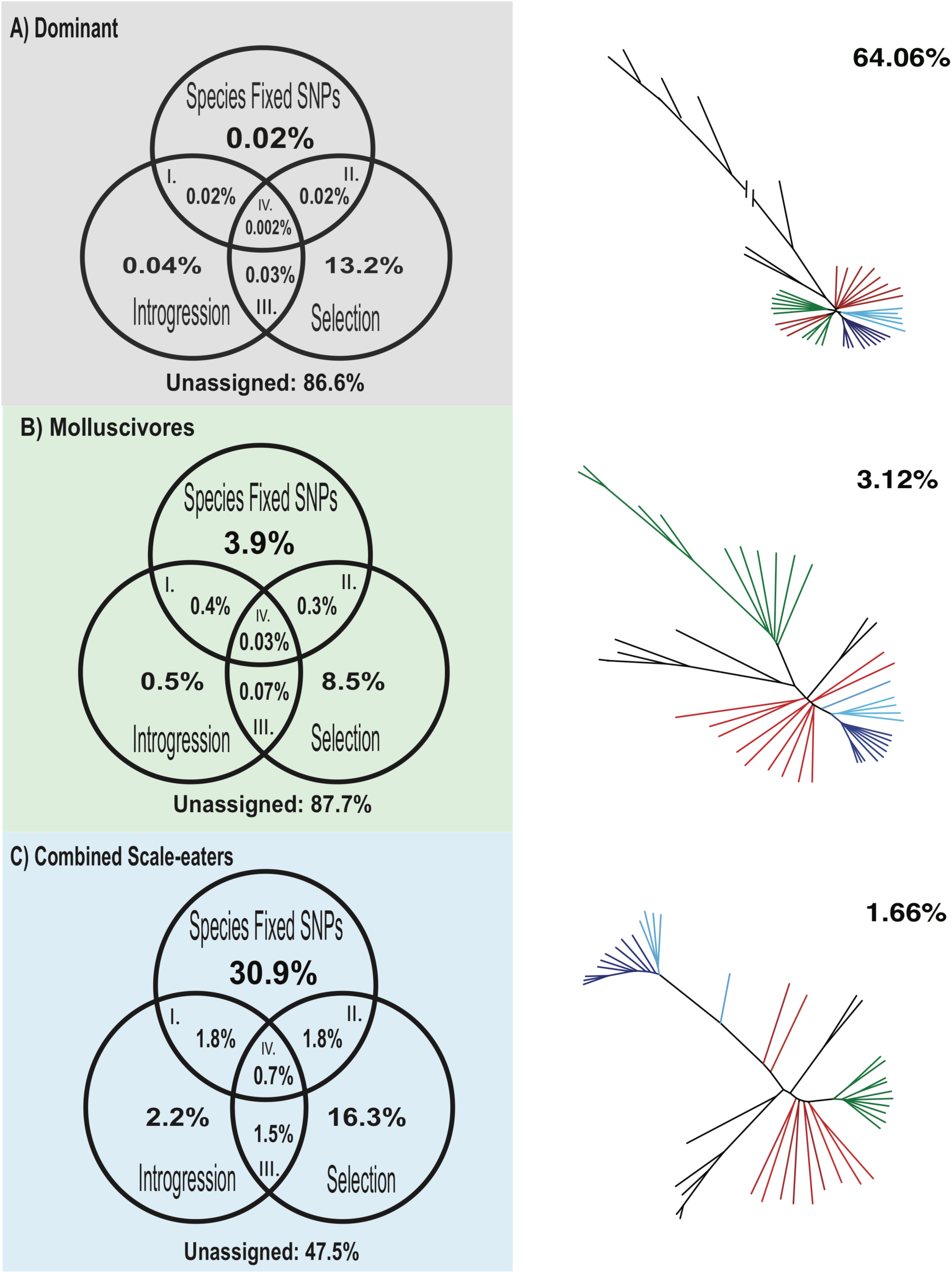
Contributions of selection and introgression to species divergence across regions assigned to different topologies. Venn diagrams of the contribution of different sources of genetic variation to speciation in this system based on fixed SNPs between the molluscivore and combined scale-eaters, significant *f_4_* values of introgression, hard selective sweeps (the lower 5% of the distribution of Tajima’s D value) in regions assigned to the (A) dominant topology, (B) molluscivore topology, and (C) combined scale-eater topology. Under each topology, we calculated the percentage of I) regions that contain introgressed genetic variation from the Caribbean contributing to species divergence, II) regions that have undergone strong selective sweeps from non-introgressed genetic variation on San Salvador, III) adaptively introgressed regions not contributing to species divergence, and IV) regions that have undergone selective sweeps of introgressed variation that contribute to species divergence of the two specialists.

### Adaptive introgression contributed to localized adaptive radiation: a case study of *ski*

In general, selective sweeps of introgressed genetic variation that contributed to species divergence between the specialists were rare. However, four of the 31 regions strongly associated with jaw size variation in the specialists [43] showed strong evidence of introgression from *Cyprinodon* species on two distant Caribbean islands (Table 1). This includes the proto-oncogene *ski,* a candidate gene with known craniofacial effects. *Ski* encodes a nuclear protein that binds to DNA and modulates transcription [51,52]. Mutations in *ski* cause marked reductions in skeletal muscle mass, depressed nasal bridges, and shortened, thick mandible bones in mice [53,54], remarkably similar to the novel craniofacial morphologies in San Salvador Island pupfishes, including increased nasal/maxillary protrusion, shortened lower jaw, and thicker dentary and articular bones [40].

**Table 1.** Adaptively introgressed 10 kb regions and gene annotations for fixed SNPs between scale-eater and molluscivore species that lie in segments assigned to the three alternative topologies. Asterisks (*) indicate SNPs in gene regions that have been associated with San Salvador pupfish jaw size variation in a previous study [43]. Bolded genes have known functional effects on craniofacial traits in a model system. Regions that are not annotated for genes are indicated with a dash (-).

| Scaffold | Position | Segment Length | $f_4$ | Fixed SNPs | Tajima's D Molluscivore | Tajima's D Scale-eater | Gene |
| --- | --- | --- | --- | --- | --- | --- | --- |
| Large-jawed scale-eater topology |  |  |  |  |  |  |  |
| KL652578.1 | 34250-35784 | 1534 | 0.34 | 7 | 0.05 | -2.16 | - |
| KL652808.1 | 48674-50395 | 1721 | -0.30 | 50 | -2.09 | -2.10 | ube2 |
| KL653236.1 | 261439-262126 | 687 | 0.27 | 11 | -2.29 | -2.11 | - |
| KL653461.1 | 31579-32075 | 496 | -0.30 | 6 | -2.02 | -2.10 | - |
| Combined scale-eater topology |  |  |  |  |  |  |  |
| KL652649.1 | 863752-863969 | 217 | 0.25 | 20 | -2.49 | -1.89 | -* |
| KL652674.1 | 781305-782379 | 1074 | 0.23 | 21 | -1.99 | -2.22 | - |
| KL652715.1 | 801693-804224 | 2531 | -0.22 | 72 | -2.19 | -1.81 | pard3* |
| KL652715.1 | 814671-815203 | 532 | -0.22 | 3 | -1.46 | -1.81 | pard3* |
| KL652761.1 | 939073-939598 | 525 | 0.29 | 7 | -1.98 | -1.81 | colgalt2 |
| KL652959.1 | 292277-293459 | 1182 | -0.25 | 5 | -1.12 | -1.81 | wnt7b |
| KL652959.1 | 294961-295670 | 709 | -0.25 | 5 | -1.12 | -1.81 | wnt7b |
| KL652983.1 | 268808-270404 | 1596 | 0.26 | 2 | -2.24 | -1.82 | **ski\*** |
| KL653171.1 | 369020-369346 | 326 | -0.25 | 2 | -2.03 | -1.89 | lbt2 |
| KL653171.1 | 370174-370702 | 528 | -0.25 | 2 | -1.11 | -1.89 | lbt2 |
| KL653356.1 | 50459-51629 | 1170 | 0.26 | 9 | -2.12 | -1.82 | srbd1 |
| KL653356.1 | 57713-59092 | 1379 | 0.26 | 9 | -2.12 | -1.82 | srbd1 |
| KL653356.1 | 75673-76452 | 779 | -0.27 | 20 | -2.21 | -1.81 | srbd1 |
| KL653706.1 | 184149-184721 | 572 | 0.36 | 31 | -2.25 | -1.89 | plekhg* |
| Molluscivore topology |  |  |  |  |  |  |  |
| KL652867.1 | 572016-572646 | 630 | -0.32 | 2 | -0.86 | -1.89 | nbea |

The adaptively introgressed segment is located at the start of *ski* in the 5’ untranslated region and contains a fixed SNP, signature of positive selection, reduced diversity in both of the specialists, and high absolute genetic divergence between the two specialists in this region compared to surrounding regions on the scaffold (Fig 5). This pattern is consistent with a hard selective sweep of an introgressed *ski* allele from outside San Salvador. To further assess the source of introgression and which specialist is the recipient, we compared F_st_ in this region between the outgroups individuals and specialists. Genetic differentiation was minimal between molluscivores and *C. laciniatus* (F_st_ = −0.15) (Fig 5) and higher in all other pairwise comparisons (F _st_ > 0.35) between the two specialists and two outgroup Caribbean pupfish species (S3 Table), indicating gene flow between the molluscivores on San Salvador Island and the generalist *C. laciniatus* on New Providence Island. Taking a closer look at the genetic variation in this region, we observe that the *ski* SNP fixed in the San Salvador molluscivores is homozygous in *C. laciniatus* and segregating in the generalists (Fig 6A), suggesting that it occurs at an appreciable frequency in the generalists. In the surrounding molluscivore genetic background of the fixed *ski* SNP is very similar to *C. laciniatus* (Fig 6B). In this region, there are only 62 variants between the molluscivores and *C. laciniatus* in our sample.

**Fig 5.**
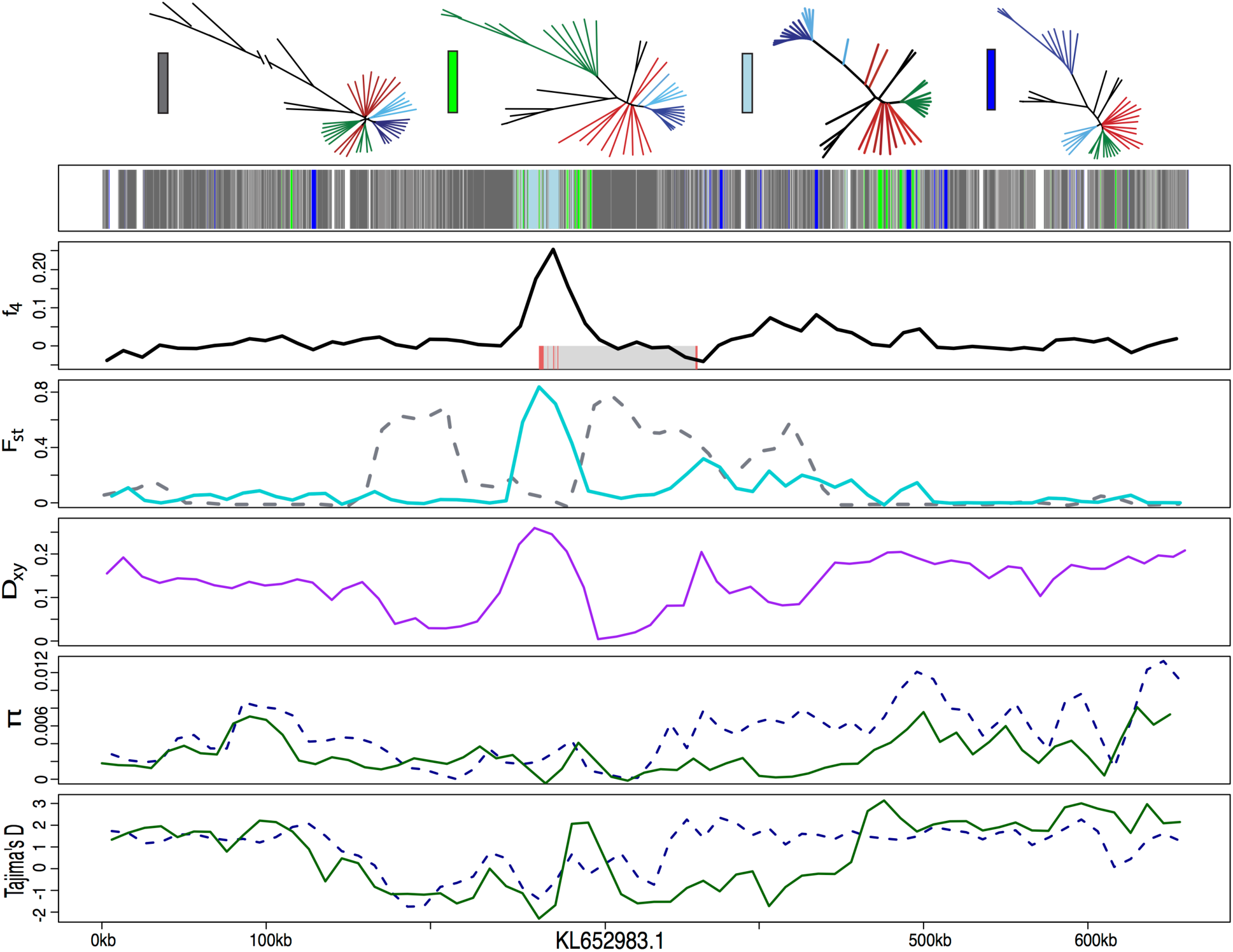
Candidate gene *ski* associated with jaw size [43] with signatures of introgression and a hard selective sweep. Row 1 shows the history assigned by SAGUARO to segments along a 600-kb scaffold (dark grey: dominant topology; blue: large-jawed scale-eater topology; light blue: combined scale-eater topology; green: molluscivore topology; light grey: all other topologies; white: unassigned segments). Row 2 shows average *f*_4_ value across non-overlapping 10-kb windows between mollsucivores/scale-eaters. Shaded grey box shows region annotated for *ski* gene with exons in red. Row 3 shows average *F_st_* value across non-overlapping 10-kb windows between molluscivores/scale-eaters (turquoise) and molluscivores/C. *laciniatus* (gray-dashed). Row 4 shows between-population divergence (*D*_xy_) across non-overlapping 10-kb windows between molluscivores/scale-eaters. Row 5 shows within-population diversity (*π*) across non-overlapping 10-kb windows (blue-dashed: scale-eater; green: molluscivore). Row 6 shows Tajima’s D across non-overlapping 10-kb windows (blue-dashed: scale-eater; green: molluscivore).

**Fig 6.**
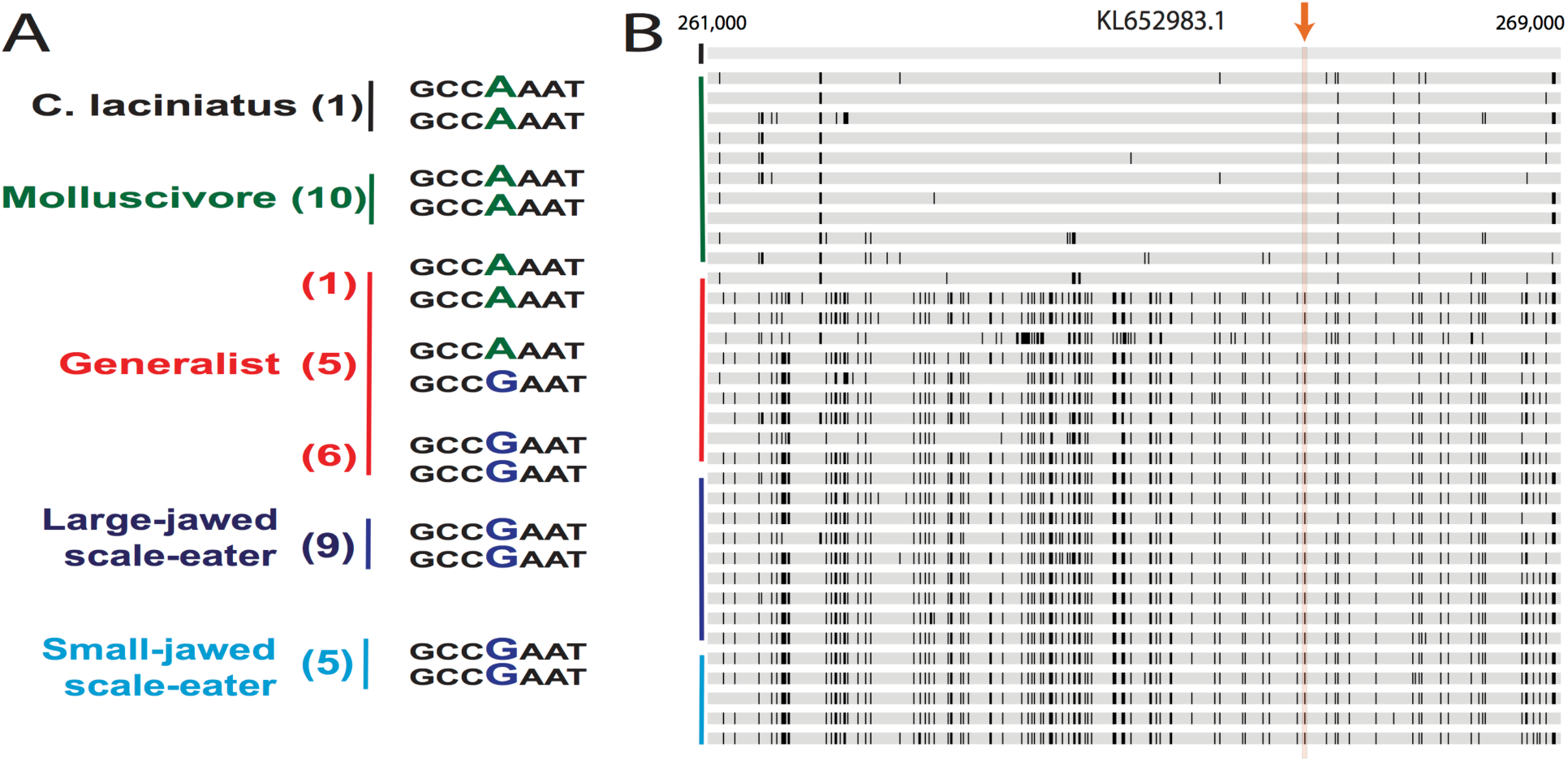
Genetic diversity surrounding the fixed variant in *ski* region assigned to the combined scale-eater topology. (A) The variant fixed between the two specialists. The number of individuals with the haplotype(s) are located in parentheses next to species names. (B) A comparison of the San Salvador genotypes (green=molluscivore; red=generalists; blue=scale-eater) with the *C. laciniatus* genotype (black) across an 8-kb window surrounding the fixed variant (orange arrow). The alleles that do not match the alleles of *C. laciniatus* are highlighted with black bars. The arrow points to the conflicting genotypes in the surrounding 8 kb region of the SNP.

We then used TREEMIX [50] to visualize the direction gene flow in adaptively introgressed regions that contributed to species divergence in the specialists. We found evidence that genetic variation from both *C. laciniatus* and *C. bondi* introgressed into both specialists (Fig 7A-D). This included the introgressed region containing *ski,* with evidence of gene flow from *C. laciniatus* into the molluscivore (Fig 7A). In other regions we found evidence of gene flow from both distant Caribbean Islands (New Providence Island, Bahamas and the Dominican Republic) into the generalist populations on San Salvador as well as gene flow from San Salvador into the outgroup Caribbean species (S4 Table).

**Fig 7.**
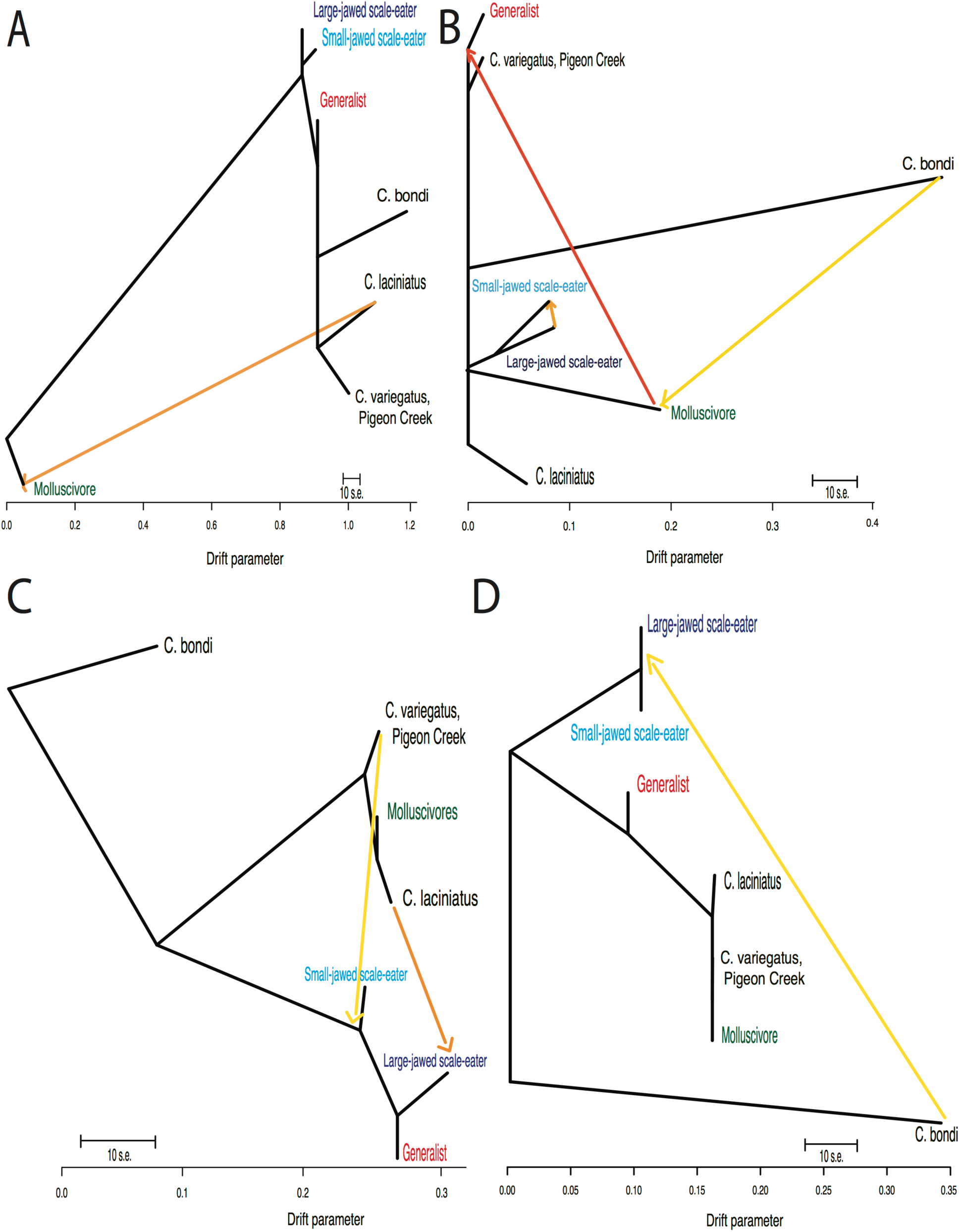
Three different histories of adaptive introgression into the molluscivore specialist from a distant Caribbean island visualized using using Treemix graphs. Adaptively introgressed region overlapping with the combined scale-eater topology supporting (A) gene flow from *C. laciniatus* into the molluscivores in the *ski* region (change in composite log-likelihood with increase in number of migration events: m=0, LnL: −320; m=1, LnL: 81), (B) gene flow from *C. bondi* into the molluscivores (m=1, LnL: 74; m=2, LnL: 107), (C) gene flow from *C. laciniatus* into the scale-eaters (m=0, LnL: −268; m=1, LnL: 95), and (D) gene flow from *C. bondi* into scale-eaters (m=2, LnL: −274; m=3, LnL: 130).

## Discussion

### Alternative topologies reveal diverse sources of genetic variation contributed to a highly localized adaptive radiation

Our investigation of genetic variation reveals that multiple sources of genetic variation from distant corners of the Caribbean have been important in the assembly of the complex phenotypes associated with the novel ecological transitions seen only on San Salvador. While species divergence appears to mostly come from selective sweeps of standing or *de novo* genetic variation (Fig 4), rare adaptive introgression has played a role in the radiation. The adaptive introgression we found in this study has come from large admixture events into San Salvador from populations as far as 742 km across the Caribbean. Our best candidate was a region containing a single fixed variant in the molluscivore specialist previously associated with jaw size variation on San Salvador containing the proto-oncogene ski, adaptively introgressed from another pupfish species on an island 300 km away (Figs 5,6, and 7A, Table 1). Importantly, our limited sampling of one individual from each of two distant islands suggests that long-distance adaptive introgression is common and arises from abundant genetic variation found in only some parts of the Caribbean.

We rarely know the source of candidate variants involved in diversification or the contributions of multiple sources of genetic variation to rapid diversification. Genomic investigations of other adaptive radiations have also inferred roles for multiple genetic sources contributing to rapid diversification. For example, in the apple maggot fly, ancient gene flow from Mexican populations introduced an inversion affecting key diapause traits that aided the sympatric host shift to apples in the United States [55]. Hybridization within Darwin’s finches also appears to play a role in the origin of new lineages through adaptive introgression of functional loci contributing to beak shape differences between species [21]. In a *Mimulus* species complex, introgression of a locus affecting flower color appears to have been a driver of adaptation in the early stages of their diversification [56]. However, even in case studies demonstrating multiple sources of genetic variation, the relative contributions to the diverse ecological traits in these radiations still remain unknown in most cases (but see [57]).

### The genomic landscape of introgression differs between sympatric trophic specialists

Only 10% of all introgressed regions in either the molluscivore or scale-eater were shared between the two. This minimal overlap may reflect the complexity of different performance demands. Performance in the two specialists involves very different sets of functional traits (i.e. higher mechanical advantage and a novel nasal protrusion in the molluscivores vs. enlarged oral jaws and adductor muscles in the scale-eaters [42]) and divergent selective regimes (narrow and shallow vs. wide and deep fitness valleys [41,58,59]). The extensive variability in the genetic variation that introgressed between the two specialists may reflect multidimensional adaptation to two distinct trophic niches in this radiation, rather than variation along a linear axis (e.g. see [60–65]).

### Did adaptive introgression trigger adaptive radiation?

Although adaptive introgression is rare and localized across the genome, it was likely important for the assembly of the complex phenotypes observed on San Salvador (e.g. *ski*). Our findings suggest that rare adaptive introgression may have been required to trigger the radiation in the presence of ecological opportunity. Indeed, a longstanding paradox in this system is why generalist populations in hypersaline lakes on neighboring islands with similar levels of ecological opportunity, lake areas, and overall genetic diversity have not radiated (Martin 2016).

The presence of introgressed adaptive variants on San Salvador could be explained through two main admixture scenarios. They could represent a large admixture event into the entire San Salvador genetic pool, a prediction of the hybrid swarm hypothesis [27], which was then sorted during the initial divergence of the specialist populations. Alternatively, the introgressed adaptive variants may represent secondary gene flow, aiding the ongoing divergence of the specialists. Our TREEMIX analyses weakly support a scenario of secondary contact for several regions, including ski, with evidence of admixture from *C. bondi* and C. *laciniatus* directly into the specialists (Fig 7). A closer investigation of the *ski* variant fixed between the specialists shows that the variant fixed in the molluscivores is segregating in the generalist population (Fig 6). This suggests that either admixture brought the adaptive *ski* allele into the generalist San Salvador population and was subsequently fixed in the molluscivores or that secondary gene flow between molluscivores and generalists has introduced the allele into the generalist through secondary contact. However, these are only exploratory inferences on directionality of gene flow and timing of introgression. They should be confirmed with demographic analyses focused on testing different scenarios of admixture into San Salvador (e.g. [26, 66–70]).

We can roughly estimate the timing of introgression for this ski region from the number of variants that have accumulated between the *C. laciniatus* and molluscivore haplotypes (n = 62 variants; Fig 6). Assuming neutrality, the observed genetic differences between the two lineages should equal 2μt the time since their divergence in each lineage and μ, the mutation rate [71]. Using mutation rate estimates ranging from 5.37×10^−7^ (phylogeny-based estimate of *Cyprinodon* substitution rate [69]) to 1.32×10^−7^ mutations site^−1^ year^−1^ (estimated from cichlid pedigree estimate of the per generation mutation rate [72] using a pupfish generation time of 6 months), introgression of the *ski* adaptive haplotype from *C. laciniatus* into the molluscivore specialist occurred between 5,700 to 23,500 years ago. The 10,000 year age estimate of the San Salvador Island radiation (based on the most recent lake drying event estimates [37–39]) falls within this window, suggesting the intriguing scenario in which widespread introgression during the last glacial maximum may have triggered adaptive radiation within pupfish populations isolated in the saline lakes of San Salvador Island during their initial formation.

While there are rare and convincing examples of hybridization leading to homoploid speciation (reviewed in [34]), no study has yet provided convincing evidence that hybridization was directly involved in triggering an adaptive radiation. For example, while there is strong evidence in Darwin’s finches that adaptive introgression of a loci controlling beak shape has contributed to phenotypic diversity of finches in the Galapagos, this hybridization occurred between members within the radiation [21]. Another recent study argued that hybridization between ancestral lineages of the Lake Victoria superflock cichlid radiations and distant riverine cichlid lineages fueled the radiations, based on evidence of equal admixture proportions across the genomes of the Victorian radiations from the riverine lineages and the presence of allelic variation in opsins in the riverine lineages which are also important in the Victoria radiation [35]. However even this fascinating finding only indirectly supports hybridization as the trigger of adaptive radiation. Allelic variation in opsins is common in most cichlid communities [73–79] and adaptive radiation is common in many African lakes [80,81]; thus it remains unclear whether the ancestral Victorian cichlid community would not have radiated without the additional influx of opsin variation from riverine lineages. Admittedly, hybridization as the necessary and sufficient trigger of adaptive radiation is a difficult prediction to test.

Those examples with more direct evidence linking hybridization to adaptation and reproductive isolation within a radiation are often special cases where a single introgressed adaptive allele automatically results in increased reproductive isolation. Examples include introgressed adaptive loci controlling wing patterns in *Heliconius* butterflies involved in mimicry and mate selection [82,83], a locus controlling copper tolerance in *Mimulus* that is tightly associated with one causing hybrid lethality [12], and loci contributing to differing insecticide resistance in the M/S mosquito mating types [84–86]. While these cases provide convincing evidence that adaptive introgression can facilitate both ecological divergence and reproductive isolation, it is still unclear whether this introgression has actually triggered or simply contributed to the ongoing process of adaptive radiation.

### A new small-jawed scale-eating species within the radiation?

We also found evidence of a distinct clade of small-jawed scale-eaters, separate from the large-jawed scale-eaters (Figs 1 and 2). The consistent clustering of this clade across the genome suggests that they are a distinct, partially reproductively isolated population on San Salvador, rather than a product of hybridization between generalists and scale-eaters in the lakes where they exist sympatrically. Indeed, the small-jawed scale-eaters appear to be more reproductively isolated than the molluscivores and generalists, as they form a distinct clade from the other species across a larger percentage of the genome than the molluscivores do (Figs 1 and 2; S1 and S2 Figs). Instead, these small-jawed scale-eaters could represent a new ecomorph that only occurs in some lake environments. They have only been observed in the five lakes connected to the Great Lake System on San Salvador (Great Lake, Mermaid’s Pond, Osprey Pond, Oyster Pond, and Stout’s Lake), but not in isolated lakes such as Crescent Pond. Consistent with this pattern of occurrence, F2 hybrid phenotypes resembling the scale-eaters have previously been shown to have extremely low survival and growth rates in these isolated lakes [58].

Small-jawed scale-eaters may also represent a stepping-stone in the evolutionary path to scale-eating. If they were a stepping stone, we might expect regions of the genome to reflect a nested relationship between the large-jawed and small-jawed scale-eaters. We see this predicted pattern in the combined scale-eater topology that underlies most of the fixed variants between the two specialists (Fig 2). Small-jawed scale-eater diets also appear to be consistent with intermediate levels of lepidophagy (scale-eating). Preliminary gut content analyses reveal that scales were found in the stomachs of 30% of small-jawed scale-eaters (*n* = 23, unpublished data J.A. McGirr) compared to 91% of large-jawed scale-eaters (*n* = 53 [42]; CHM unpublished data).

### The complex evolutionary history of the San Salvador pupfish radiation illustrates the inadequacy of the “species tree” view

Localized regions containing traits relevant to understanding speciation in this system would be obscured by a genome-wide species tree summary. Our dominant history contained only 10% of the variants fixed between species. Furthermore, this dominant history is not a genome-wide summary of coalescence akin to a species tree. Instead, it provides a meaningful segmentation of the genome into speciation-relevant and largely geography-relevant histories [87,88]. Speciation genomic studies of many other radiations have similarly found evidence of localized regions that are highly differentiated relative to the rest of the genome, demonstrating that specific regions play an important role in maintaining reproductive isolation in the face of gene flow [21, 24, 89–92]. Understanding rapid adaptive radiation requires a new framework, one built upon the fact that different regions of the genome correspond to different evolutionary histories, in order to understand the mechanisms that drive diversification in this system. Within this framework, species trees and other genome-wide summaries of evolutionary history are meaningless, or worse, misleading for understanding what triggers adaptive radiations (see also [89,93,94]).

### Conclusion

Here we demonstrate that the complex phenotypes associated with the novel ecological transitions seen on San Salvador arose from multiple sources of genetic variation spread across the Caribbean. The variation important to this radiation is localized in small regions across the genome that are obscured by genome-wide summaries of the history of the radiation. Species divergence appears to mostly come from selective sweeps of standing or *de novo* genetic variation on San Salvador, but rare adaptive introgression events may also be necessary for the radiation. This genomic landscape of introgression is variable between the specialists and has come from large admixture events from populations as far as 742 km across the Caribbean. Our findings that multiple sources of genetic variation contribute to the San Salvador radiation suggests a complex suite of factors that include rare adaptive introgression, may be required to trigger adaptive radiation in the presence of ecological opportunity.

## Methods

### Study system and samples

Individual pupfish were caught in hypersaline lakes on San Salvador Island in the Bahamas with either a hand or seine net in 2011, 2013, and 2015. Samples were collected from seven isolated lakes on this island (Crescent Pond, Great Lake, Little Lake, Mermaid’s Pond, Moon Rock Pond, Oyster Lake, and Stout’s Lake) and one estuary (Pigeon Creek). 13 *Cyprinodon variegatus* were sampled from all eight lakes on San Salvador Island; 10 *C. brontotheroides* were sampled from four lakes; and 14 *C. desquamator* were sampled from six lakes as the specialist species occur in sympatry with the generalists in only some of the lakes. Individual pupfish that were collected from other localities outside of San Salvador Island served as outgroups to the San Salvador Island radiation, including closely related *C. laciniatus* from Lake Cunningham, New Providence Island in the Bahamas, *C. bondi* from Etang Saumautre lake in the Dominican Republic, *C. diabolis* from Devil’s Hole in California (collected as a dead specimen by National Park Staff in 2012), as well as captive-bred individuals of *C. simus* and *C. maya* originating from Laguna Chicancanab, Quintana Roo, Mexico. Fish were euthanized by an overdose of buffered MS-222 (Finquel, Inc.) following approved protocols from University of California, Davis Institutional Animal Care and Use Committee (#17455) and University of California, Berkeley Animal Care and Use Committee (AUP-2015-01-7053) and stored in 95-100% ethanol. Only degraded tissue was available for *C. diabolis,* as described in [69].

### Genomic sequencing and bioinformatics

DNA was extracted from muscle tissue using DNeasy Blood and Tissue kits (Qiagen, Inc.) and quantified on a Quibit 3.0 fluorometer (Thermofisher Scientific, Inc.). PCR-free Truseq-type genomic libraries were prepared using the automated Apollo 324 system (WaterGen Biosystems, Inc.) at the Vincent J. Coates Genomic Sequencing Center (QB3). Samples were fragmented using Covaris sonication, barcoded with Illumina indices, and quality checked using a Fragment Analyzer (Advanced Analytical Technologies, Inc.). Nine to ten samples were pooled in four different libraries for sequencing on four lanes of Illumina 150PE Hiseq4000.

426 million raw reads were mapped from 42 individuals to the *Cyprinodon* reference genome (NCBI, *C. variegatus* Annotation Release 100, total sequence length = 1,035,184,475; number of scaffold = 9,259, scaffold N50, = 835,301; contig N50 = 20,803) with the Burrows-Wheeler Alignment Tool [95] (v 0.7.12). Duplicate reads were identified using MarkDuplicates and BAM indices were created using BuildBamIndex in the Picard software package (http://picard.sourceforge.net(v.2.0.1)). We followed the best practices guide recommended in Genome Analysis Toolkit [96](v 3.5) to call and refine our SNP variant dataset using the program HaplotypeCaller. Since we lacked high-quality known variants that are typically required as a reference to filter SNP variants, we filtered SNPs based on the recommended hard filter criteria (i.e. QD < 2.0; FS < 60; MQRankSum < −12.5; ReadPosRankSum < −8) [84,96]. Our final dataset after filtering contained 16 million variants and a mean sequencing coverage of 7.2X per individual (range: 5.2–9.3X).

### Characterization of genomic heterogeneity in evolutionary relationships across individuals

We used the machine learning program SAGUARO [44] to identify regions of the genome that contain different signals about the evolutionary relationships across San Salvador Island and outgroup *Cyprinodon* species. Saguaro combines a hidden Markov model with a self-organizing map to characterize local phylogenetic relationships among aligned individuals without requiring *a priori* hypotheses about the relationships. This method infers local relationships among individuals in the form of genetic distance matrices and assigns segments across the genomes to these topologies. These genetic distance matrices can then be transformed into neighborhood joining trees to visualize patterns of evolutionary relatedness across the genome. Three independent runs of SAGUARO were started using the program’s default settings and each was allowed to assign 15 different histories across the genome. The 15^th^ history and additional histories that we investigated tended to be uninformative about the evolutionary relationships among individuals and represented less than 0.5% of the genome. To determine how many histories to estimate, analogous to a scree plot [97,98], we plotted the proportion of the genome explained by a hypothesized history and looked for an inflection point in the section of the plot where hypothesized histories explained little of the genome. We also looked at the neighborhood joining trees to assess whether additional histories were informative about the evolutionary relationships among individuals (S9 Fig). We excluded the last history (15^th^) from downstream analyses due to lack of genetic distinction at both the level of populations and species included in the proposed genetic distance matrix and the low percentage of the genome assigned to it. The 14 histories included and the total percentages of the genome assigned to them were robust across all three independent runs.

### Characterization of introgression patterns across the genome

We characterized the heterogeneity in introgression across the genome using *f_4_* statistics that were initially developed to test for introgression among human populations [48–50]. The *f_4_* statistic tests if branches among a four-taxon tree lack residual genotypic covariance (as expected in the presence of incomplete lineage sorting and no introgression) by comparing allele frequencies among the three possible unrooted trees. A previous study [41] provides evidence of potential admixture with the Caribbean outgroups species used in this study, preventing their use in a D-statistic framework which requires explicit outgroup designation.

To look for evidence of gene flow across the Caribbean, we focused on tests of introgression with the two outgroup clades from our sample that came from other Caribbean islands in the Bahamas and Dominican Republic. Based on the tree ((P1, P2),(C. *laciniatus, C. bondi)), f_4_* statistics were calculated for all three possible combinations of P1,P2 among the pooled populations of generalists, scale-eaters, and molluscivores on San Salvador Island. These *f_4_* statistics were calculated using the population allele frequencies of biallelic SNPs and summarized over windows of 10 kb with a minimum of 50 variant sites using a custom python script (modified from ABBABABA.py created by Simon H. Martin, available on https://github.com/simonhmartin/genomics_general), allowing for up to 10% missing data within a population per site. All 10 and 14 individuals from San Salvador were used in the tests for the molluscivores and scale-eaters, respectively, for the comparison to the molluscivore and combined scale-eater histories. In another calculation of *f_4_* statistics across the genome, the 5 small-jawed scale-eater individuals were excluded for the comparison to the large-jawed scale-eater topology. Although only single individuals from New Providence Island, Bahamas and the Dominican Republic were used to represent *C. laciniatus* and *C. bondi* in the *f_4_* tests, these individuals that were sequenced are a random sample from the populations and should be representative. This resulted in 100,276 *f_4_* statistics (mean *f_4_* = −2×10^−4^) calculated across the genome for the test that included all scale-eaters and 100,097 *f_4_* statistics (mean *f_4_* =-9×10^−5^) for the test excluding the small-jawed scale-eaters. The significance of *f_4_* values was assessed through the calculation of Z-scores using the mean and standard deviation of *f_4_* values calculated from all regions across the genome. These Z-scores were then transformed into P-values and significance was assessed using a Bonferroni-corrected significance threshold of 5×10^−7^.

It is difficult to distinguish between genetic variation that is similar among taxa due to introgression from a hybridization event and that from ancestral population structure, so some of the regions with significant *f_4_* values may represent the biased assortment of genetic variation into modern lineages from a structured ancestral population. A recent simulation study [99] found that extending the use of genome-wide introgression statistics such as Patterson’s D statistic and *f_4_* statistics to small genomic regions can result in a bias of detecting statistical outliers mostly in genomic regions of reduced diversity. We assessed whether our identified regions of introgression were predominately from regions of low diversity by comparing *D_xy_* and π estimates across the detected regions of introgression in comparison to the genome-wide estimates.

To visualize gene flow among the Caribbean populations included in this study, we used TREEMIX v1.12 [50] to estimate population graphs with 0-4 admixture events connecting populations. Population graphs were estimated for each region with a significant *f_4_* value, each with a minimum of 50 SNPs and a block size of 1. The number of admixture events was estimated by comparing the rate of change in log likelihood of each additional event, an approach similar to one used in Evanno et al. ([100]; also see [41]). However, this analysis should be viewed only as an exploratory tool as the reliability of TREEMIX to detect the number of admixture events has not been tested. This analysis was designed to be applied on genome-wide allele frequencies and estimates covariance in allele frequencies among populations in branch lengths using a model that assumes allele frequency differences between populations are solely caused by genetic drift [50]. The use of fewer SNPs (≥50) in our window-based approach also makes it harder to reliably distinguish between the different likelihoods for the number of migration events. The reliability of inference under these conditions has not been evaluated, however the migration events inferred in our TREEMIX results were consistent with our findings from our formal *f_4_* test for gene flow.

### Comparison of patterns of introgression to patterns of genetic divergence and diversity

We then calculated several population genetic summary statistics in sliding windows across the genome to compare to the *f*4 patterns of introgression: F_st_, within-population nucleotide diversity (π) for pairwise species comparisons, and Tajima’s D estimates of selection in each species. These statistics were calculated in non-overlapping sliding windows of 10 kb using ‘wier-fst-pop’, ‘window-pi’, and ‘TajimaD’ functions in VCFtools v.0.1.14 [101]. Negative values of Tajima’s D indicate a reduction in nucleotide variation across segregating sites [102], which may result from hard selective sweeps due to positive selection. We took the regions with Tajima’s D values in the 95% percentile from the lower tail of the distribution as regions under strong positive selection. Population genetic divergence (D_xy_) between molluscivores and scale-eaters was calculated over the same 10-kb windows of the *f_4_* tests using the python script popGenWindows.py created by Simon Martin (available on https://github.com/simonhmartin/genomics_general). Within regions of overlap between significant *f_4_* values, strongly negative Tajima’s D values, fixed SNPs between the two specialists, and alternative topologies, we looked for annotated genes and searched their gene ontology in the phenotype database ‘Phenoscape’ [103–106] and AmiGO2 [107] for skeletal system effects. Skeletal features, particularly craniofacial morphologies such as jaw length, have extremely high rates of diversification among the species on San Salvador Island [36,41] and likely play a key role in the diversification of this group.

## Data accessibility

All datasets used for this study will be deposited in Dryad and the NCBI Short Read Archive.

## Acknowledgements

This study was funded by a Daphne and Ted Pengelley Award from the Center for Population Biology and the University of North Carolina to CHM. We thank J. McGirr, J. Poelstra, and the UNC EEGAS journal club for helpful discussion of this manuscript; K. Gould for performing DNA extractions; M. Grabherr for discussion of filtering strategies; members of the American Killifish Association for supplying tissues, in particular A. Morales, M. Schneider, J. Cokendolpher, and A. Kodric-Brown; R. Hanna, K. Guerrero, A. Valdes Gonzalez, and L. Simons for assistance obtaining permits; the Gerace Research Centre for accommodation; the governments of the Bahamas, Dominican Republic, and the National Park Service and U.S. Fish and Wildlife Service for permission to collect pupfish samples; the Vincent J. Coates Genomic Sequencing Center and Functional Genomics Laboratory, supported by NIH S10 OD018174 Instrumentation Grant, for performing whole-genome sequencing; and the UNC ITS Research Computing Services for computational resources. We declare no competing financial interests.

## Author Contributions

EJR wrote the manuscript and conducted all analyses. CHM collected the samples and provided the genomic data. Both authors contributed to the development of ideas presented in the study and revised the manuscript.

## Supporting Information

**S1 Fig. Hypothesized histories featuring a monophyletic San Salvador clade and the percent of the genome assigned to the histories.** Black lineages are the *Cyprinodon* outgroups, red lineages are the San Salvador generalists, green lineages are the San Salvador molluscivores, dark blue lineages are the large jawed scale-eaters and light blue lineages are the small-jawed scale-eaters.

**S2 Fig. Hypothesized histories featuring a non-monophyletic San Salvador clade and the percent of the genome assigned to the histories.** Black lineages are the *Cyprinodon* outgroups, red lineages are the San Salvador generalists, green lineages are the San Salvador molluscivores, dark blue lineages are the large jawed scale-eaters and light blue lineages are the small jawed scale-eater.

**S3 Fig. Visualization of introgression across the genomes of molluscivores and scale-eaters.** Manhattan plot of the *f_4_* values between the San Salvador molluscivores, scale-eaters, *C. laciniatus* from New Providence Island, Bahamas and *C. bondi* from Etang Saumautre, Dominican Republic. Alternating gray/black colors indicate different scaffolds, starting with the largest scaffolds in the top row and the smallest scaffolds in the bottom row. Dotted red lines mark the of Bonferroni-corrected significance level for the *f_4_* values (P-value < 5.2×10^−7^).

**S4 Fig. Visualization of introgression across the genomes of molluscivores and generalists.** Manhattan plot of the *f*_4_ values between the San Salvador molluscivores, generalists, *C. laciniatus* from New Providence Island, Bahamas and *C. bondi* from Etang Saumautre, Dominican Republic. Alternating gray/black colors indicate different scaffolds, starting with the largest scaffolds in the top row and the smallest scaffolds in the bottom row.

**S5 Fig. Visualization of introgression across the genomes of scale-eaters and generalists.** Manhattan plot of the *f*_4_ values between the San Salvador large-jawed scale-eaters, small-jawed scale-eaters, *C. laciniatus* from New Providence Island, Bahamas and *C. bondi* from Etang Saumautre, Dominican Republic. Alternating gray/black colors indicate different scaffolds, starting with the largest scaffolds in the top row and the smallest scaffolds in the bottom row.

**S6 Fig. Comparison of *f_4_* to genetic diversity statistics over 10-kb non-overlapping windows.** Red dots indicate 10-kb regions with signals of introgression above Bonferroni-corrected significance level the *f_4_* values (P-value < 5.2×10^−7^). The *f_4_* statistic of a region compared to A) *F_st_* between molluscivores and scale-eaters in the region, B) *D_xy_* between molluscivores and scale-eaters on San Salvador, C) within-population diversity in molluscivores, and D) within-population diversity in San Salvador scale-eaters.

**S7 Fig. Comparison of f4 to genetic diversity statistics over 10-kb non-overlapping windows.** Red dots indicate 10-kb regions with signals of introgression above Bonferroni-corrected significance level the f_4_ values (*P*-value < 5.2×10^−7^). The f4 statistic of a region compared to A) Fst between molluscivores and scale-eaters in the region, B) D_xy_ between molluscivores and scale-eaters on San Salvador, C) within-population diversity in molluscivores, and D) within-population diversity in San Salvador scale-eaters.

**S8 Fig. The percentage of segments assigned to large-jawed scale-eater topology that contain signatures of species divergence, selection, and introgression.** Venn diagrams of the contribution of different sources of genetic variation to speciation in this system based on the overlap of regions with fixed SNPs between the molluscivore and large-jawed scale-eater, significant *f_4_* values of introgression, the lower 5% of the distribution of Tajima’s D value with positive selection and their overlap. Under each topology, we calculated the percentage of I) regions that contain introgressed genetic variation from the Caribbean contributing to species divergence, II) regions that have undergone strong selective sweeps from non-introgressed genetic variation on San Salvador, III) adaptively introgressed regions not contributing to species divergence, and IV) regions that have undergone selective sweeps of introgressed variation that contribute to species divergence of the two specialists.

**S9 Fig. The proportion of the genome assigned to each topology by SAGRUARO.** The insert is a closer look at the 13 topologies assigned to the smallest proportion of the genome and the uninformative 15^th^ topology hypothesized. This shows a saturation in the variation in topologies across the genome at 14.

**S1 Table: Hypothesized topologies from the SAGUARO analysis.**

**S2 Table: Summary statistics in adaptively introgressed regions.**

**S3 Table: Pairwise genetic divergence (Fst) between molluscivores, scale-eaters, *C. laciniatus* and *C. bondi.***

**S4 Table: Summary of admixture events inferred by TREEMIX for the 21 adaptively introgressed regions assigned to the three alternative topologies**. San Salvador Island generalist (G), San Salvador Island large-jawed scale-eater (L), San Salvador Island small-jawed scale-eater (S), San Salvador Island molluscivore (M), *C laciniatus* from New Providence Island Bahamas (CUN), *C. bondi* from Dominican Republic (ETA), most recent common ancestor of Caribbean pupfish lineages (MRC).

